# Extending Conformational Ensemble Prediction to Multidomain Proteins and Protein Complex

**DOI:** 10.64898/2026.01.14.699584

**Authors:** Junjie Zhu, Sören von Bülow, Hongyi Liu, Kresten Lindorff-Larsen, Hai-Feng Chen

## Abstract

Proteins execute cellular functions through a structural continuum ranging from stable, folded domains to highly dynamic intrinsically disordered regions (IDRs). Conformational ensembles represent the set of three-dimensional structures a protein adopts under a specific set of conditions, and underlie essential processes from catalysis to complex signaling networks. While deep learning has revolutionized structure prediction, capturing the distinct conformational diversity of folded and disordered regions—especially within multidomain proteins and large assemblies—remains a fundamental challenge. Here we introduce IDPFold2, a generative framework that models the heterogenous protein thermodynamics by integrating a Mixture-of-Experts architecture into the flow matching framework. By routing residues from different regions to specialized expert networks, IDPFold2 accurately predicts conformational ensembles for folded domains, IDRs and multidomain proteins. IDPFold2 outperforms state-of-the-art methods in capturing key functional states and fitting the experimental observations across local and global scales. Furthermore, we describe an extension of IDPFold2 to protein assemblies, deciphering the complex binding modes of IDRs within large macromolecular complexes, providing a generalizable tool for exploring the dynamic proteome.

## Introduction

Proteins are fundamental biomolecules that execute vast cellular functions. The remarkable sequence variability and conformational flexibility inherent to proteins enable cells to mount targeted and rapid responses to both internal signals and external stimuli^1,2^. A central goal in molecular biology has been to gain mechanistic insights into how proteins adopt diverse structures and how this flexibility governs molecular recognition and cellular responses^3,4^.

Protein systems can be broadly categorized into folded domains and intrinsically disordered regions (IDRs) based on their inherent dynamic behavior (Fig. 1A)^5^. While the structure-function paradigm has traditionally focused on folded domains with limited structural fluctuations, disordered residues constitute ca. one third of the human proteome, and most human proteins contain both ordered and disordered regions^6,7^. The low energy barrier and large solvated area enable IDRs to serve as critical regulatory hubs, mediating long-range allostery and promiscuous binding interactions essential for signal transduction^8–10^. Consequently, deciphering the interplay between folded domains and IDRs is a prerequisite for mechanistic understanding of cellular physiology and the development of therapeutics against complicated targets^11^.

**Figure 1.**
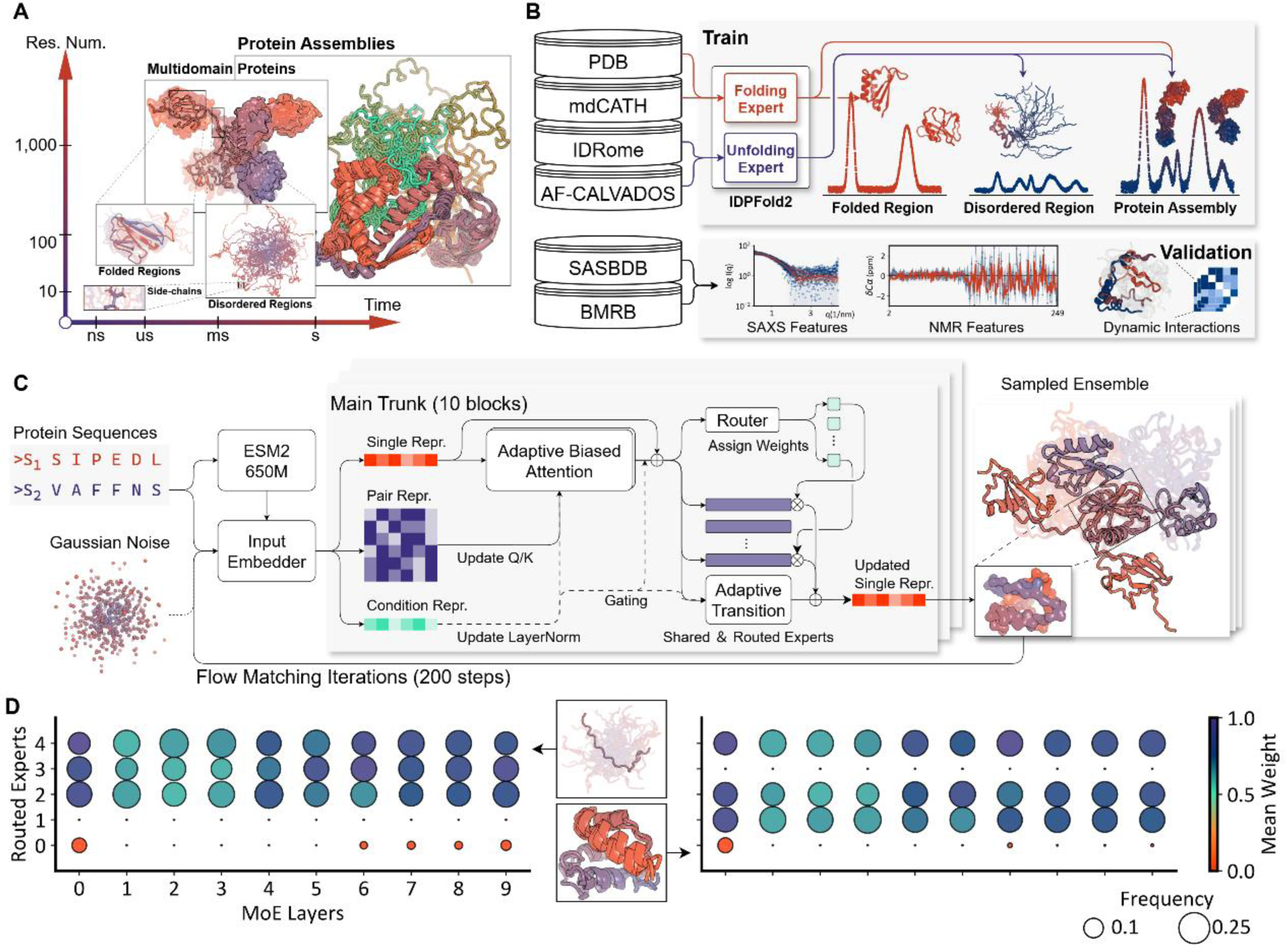
Overview of the IDPFold2 framework. (A) Proteins exhibit dynamic behaviors across a broad timescale continuum. (B) Data integration strategy. IDPFold2 is trained on static structures (PDB) and simulation trajectories covering folded domains, IDRs, MDPs and multimers. Experimental data from SASBDB and BMRB are collected for evaluation. (C) Network architecture. (D) Expert utilization analysis. Activation patterns for disordered peptide Hst5 (right) and folded domain 1w0tA00 (left) reveal that distinct experts are recruited for different structural topologies.

Despite the resolution of over 200,000 static structures, capturing the full dynamic spectrum of proteins using experimental techniques remains a formidable challenge^12^. Nuclear Magnetic Resonance (NMR), Small Angle X-ray Scattering (SAXS), and Fluorescence Resonance Energy Transfer (FRET) provide essential insights into flexibility in solution but are often limited by low resolution or the difficulty to deconvolute complex dynamic ensembles^13–15^. While Cryogenic Electron Microscopy (Cryo-EM) has revolutionized structural biology, resolving highly heterogeneous regions like IDRs from 2D density maps remains notoriously difficult^16^.

Computational approaches have long sought to bridge this gap. Molecular Dynamics (MD) simulations generate conformational ensembles by updating the velocity and position of target molecule with a predefined force field iteratively. Early iterations of all-atom protein force fields were predominantly tested on folded proteins and are readily accessible for simulating their limited dynamics^17^. More recently, various force fields have been re-parameterized for IDRs, showing better alignment with experimental observations for these highly dynamic proteins^18–21^.

Recently, deep learning has emerged as a powerful alternative. Compared to traditional MD simulations, generative models offer the advantage of generating greater conformational diversity at significantly lower computational costs and increased scale. Distributional Graphormer (DiG) and AlphaFlow represent early investigation into modeling the motions of folded protein systems^22,23^. BioEmu, trained on a large-scale dataset, can predict various motions and folding energies of folded proteins but still struggles to predict the global features of IDRs accurately^24^. Meanwhile, a series of works including IDPFold, IDPForge and IDP-o represent efforts in modeling the thermodynamics of IDRs, achieving results comparable to MD simulations in fitting ensemble-averaged features across both local and global scales^25–27^.

Despite these advances, a major challenge in modeling protein ensembles lies in distinguishing and accurately handling the distinct conformational diversity of folded regions and IDRs simultaneously. In multidomain proteins (MDPs) and protein assemblies where folded regions and IDRs coexist, methods optimized for folded proteins fail to sample the diverse conformations mediated by IDRs, while IDR-specific methods struggle to maintain the stability and native secondary structures within folded domains^28,29^. Failure to model both components hinder the broader application of these predictive methods.

Recent computational strategies have attempted to bridge this gap by explicitly compartmentalizing dynamics. CALVADOS 3 employs center-of-mass representations with elastic network constraints to preserve folded domains, while methods like AFflecto and IDPForge restrict conformational sampling to disordered regions, treating folded segments as rigid bodies and requiring manual masking^26,28,30^. bAIes statistically combine random coil models with AlphaFold2 predictions in a Bayesian network to produce ensembles^31^. However, these workflows typically rely on a priori knowledge of domain boundaries to enforce domain stability. By imposing artificial rigidity or requiring user-defined constraints, they often preclude the local fluctuations within domains and the coupled folding-binding events that define the function of complex assemblies.

Here, we developed IDPFold2, a generative deep-learning system that accurately models the thermodynamics of folded and disordered regions within a single framework. By integrating Mixture-of-Experts (MoE) into the diffusion module of AlphaFold3, IDPFold2 routes residues from distinct regions to specialized expert networks. Trained via flow matching, IDPFold2 enables the accurate and rapid generation of conformational ensembles for folded domains, IDRs, and MDPs solely from sequence. We demonstrate that IDPFold2 outperforms state-of-the-art methods on a comprehensive benchmark of 439 protein with SAXS data and 659 proteins with NMR chemical shifts, providing unprecedented accuracy across local and global scales. Furthermore, by assembling protein multimers during training, we expanded the application of IDPFold2 to predicting ensembles of protein assemblies. IDPFold2 successfully deciphers the binding modes of IDRs within large protein assemblies, offering a new lens into the dynamic molecular sociology of the cell.

## Results

### Overview

We present IDPFold2, a generative deep-learning framework designed to capture the multi-scale thermodynamics of complex protein systems. To address the scarcity of experimental data for protein assemblies, we trained the model on a composite dataset that integrated static single-chain structures from the Protein Data Bank (PDB) with extensive all-atom and coarse-grained simulation trajectories covering folded domains, IDRs, and MDPs (Fig. 1B). By reassembling static single-chain structures into multimers during training, the model learns to infer inter-chain interactions and steric constraints that are typically absent from single-chain ensemble datasets.

To overcome the intrinsic disparity between the amplitudes and types of motions in folded domains and IDRs, IDPFold2 incorporates a Mixture-of-Experts (MoE) architecture. Building on the diffusion module of AlphaFold3, we replaced the standard transition layers with sparse MoE layers, facilitating the context-dependent routing of structural tokens to specialized expert networks (Fig. 1C). IDPFold2 operates on Cα traces rather than all-atom coordinates to reduce computational overhead and maximize the diversity of sampled ensembles. Built within a flow-matching framework, IDPFold2 iteratively denoises structures from a Gaussian distribution, conditioned on sequence features extracted by ESM2-650M.

We validated the MoE hypothesis by analyzing expert utilization during the generation of the disordered peptide Hst5 and the folded domain 1w0tA00 (Fig. 1D). Non-uniform activation patterns were observed for both proteins, and specific experts were preferentially recruited for disordered regions versus folded topologies. This differential activation confirms that the router layers discriminate between structural classes, enabling the specialized processing required to handle the hierarchical thermodynamics of complex protein systems.

### Predicting global compaction across the order-disorder continuum

We validated IDPFold2 on a test set of 1,258 single-chain proteins, spanning the continuum from globular folded states to fully expanded coils. The test set consists of structural references from PDB, MD simulations, and experimental data from SASBDB and BMRB. IDPFold2 recapitulated the polymer scaling laws governing protein dimensions, sampling ensembles that span the continuum from globular folded states to fully expanded coils (Fig. 2A).

**Figure 2.**
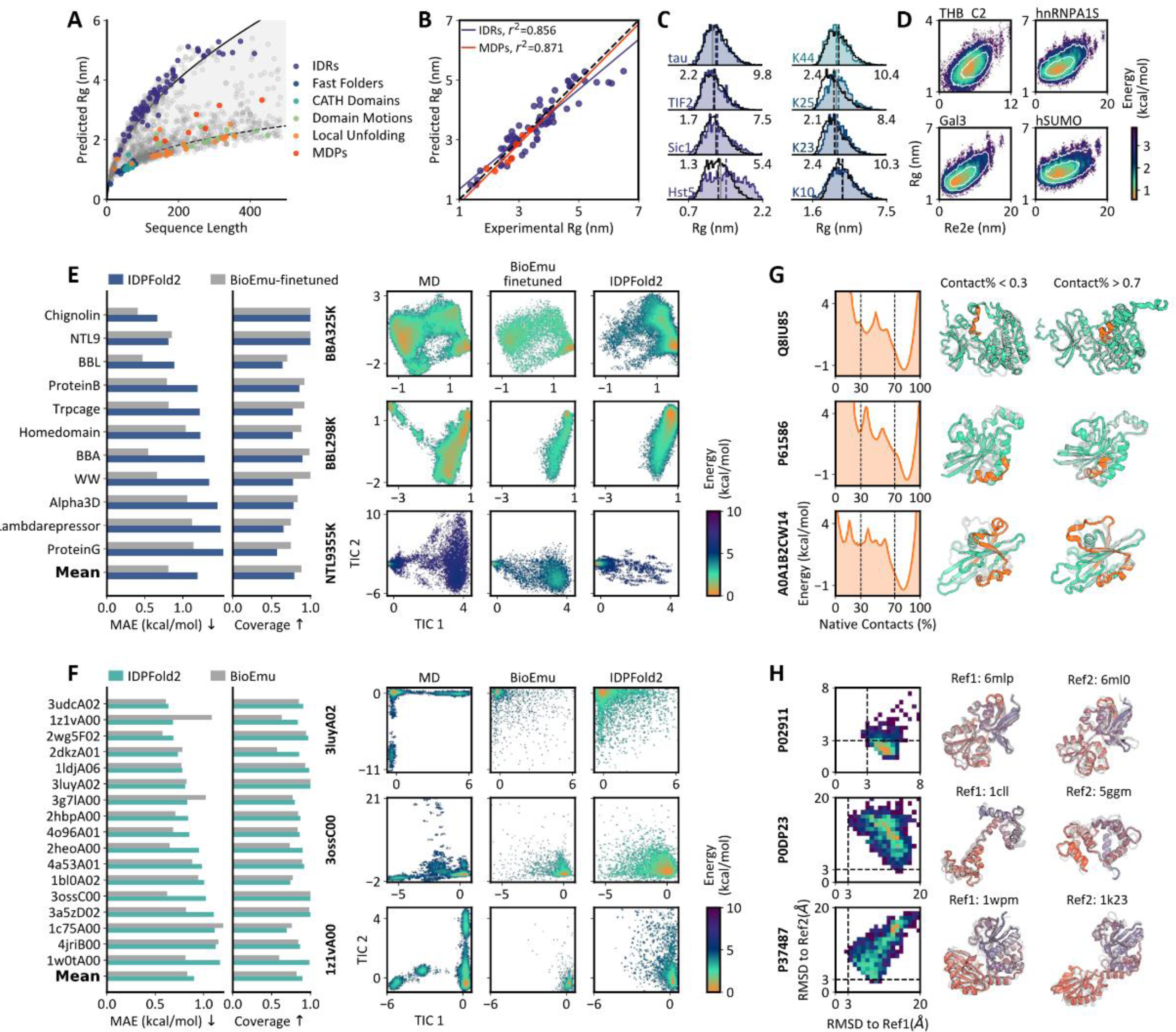
Prediction of ensembles across the order-disorder continuum. (A) Predicted ensemble-averaged Rg for 1,258 test proteins. (B) Correlation between predicted and experimental Rgs for 73 IDRs and 7 MDPs. (C) Rg distributions for 8 IDRs comparing IDPFold2 (colored) with CALVADOS 3 simulations (black). Grey vertical lines denote experimental Rgs. (D) Coupled Rg vs. End-to-End distance (Re2e) distributions for 4 MDPs. Simulation distributions are showed as white contours. (E) DESRES fast-folding proteins. Left: MAE of free-energy differences. Middle: landscape coverage. Right: TIC landscape projection. (F) CATH domains (metrics same as in E). (G) Local unfolding. Left: Fraction of native contacts. Right: locally folded and unfolded structures generated by IDPFold2. Regions undergoing local transitions are colored in orange. (H) Domain motions: Left: RMSD to reference PDB structures. Right: generated structures capturing open/closed states. Reference structures are colored in grey.

For a subset of 73 IDRs and 7 MDPs derived from CALVADOS 3, IDPFold2 predictions maintained a strong linear correlation with experimental radii of gyration (Rg), exhibiting low relative error (Fig. 2B)^28^. Predicted ensembles matched the accuracy of established coarse-grained simulations by CALVADOS3 and outperformed other deep learning approaches, demonstrating the ability of IDPFold2 in capturing IDR expansion and maintaining domain poses in MDPs (Fig. S1).

Beyond ensemble-averaged property, IDPFold2 captured the underlying distributional heterogeneity of conformational ensembles. It reproduced the broad Rg distributions on IDRs, and captures the coupled distributions of Rg and end-to-end distance (Re2e) on MDPs observed in simulations (Fig. 2C, D; Fig. S2). These distributions indicated that the model effectively sampled diverse domain arrangements instead of merely predicting correct average sizes, which is a critical feature often missed by models trained exclusively on folded structures.

### Sampling conformational changes in folded domains

Beyond global compaction, we assessed the model’s ability to capture fine-grained meta-stabilities in folded domains using the BioEmu benchmark^24^. For 12 fast-folding proteins, we used the total 8.2ms simulations by Lindorff-Larsen *et al.* as reference^32^. Ensembles were projected onto the two slowest time-independent components (TICs) to construct free-energy landscapes. Coverage of sampled range on the landscape and mean absolute error (MAE) of free-energy differences were calculated, reflecting the diversity and accuracy of generated ensemble distribution respectively. IDPFold2 demonstrated robust zero-shot generalization, reaching coverage and MAE close to BioEmu finetuned by leave-one-out cross-validation strategy (Fig. 2E, Fig. S3).

In stable CATH domains where simulations were run for 100μs per system, IDPFold2 achieved higher landscape coverage than BioEmu (Fig. 2F). Given that the reference MD simulations were shorter than those for the fast-folding proteins and were performed at 300K, unfolding events are rarely sampled in the trajectories. IDPFold2 explored a broader conformational space, including accessible partially unfolded states that were rarely sampled in simulations or by BioEmu (Fig. S4). Meanwhile, IDPFold2 maintains the native basin accurately with a similar MAE to BioEmu, offering a comprehensive view of the equilibrium distribution and accessible unfolded states.

We further considered the local unfolding process in larger protein systems with experiment-resolved key conformations. The fraction of native contacts was reported where the fraction > 0.7 could typically denote a well-folded structure and fraction < 0.3 was locally unfolded (Fig. 2G). The calmodulin binding motif in CaM kinase ID (Q8IU85) is autoinhibited in basal state by a neighbor regulatory domain, organizing a short helix that does not access substrates^33^. Binding of calmodulin pulls the regulatory domain away, and exposes the binding motif as a loop to ensure sufficient binding to this hydrophobic motif. IDPFold2 samples the full range from folded to unfolded state for this motif. Specifically, a major of basal state was sampled while unfolded state were relatively rare, corresponded exactly to the higher stability of basal state at no substate condition. Similar results were observed in the switch II region of transforming protein RhoA (P61586), where a helix is organized and pulled towards target nucleotide at GDP-bound state (native) and unfold at absence of GDP^34^. IDPFold2 also samples the distinct conformations of AvrM14-B (A0A1B2CW14) where the C-terminal domain swaps in forming dimer^35^. IDPFold2 predicts the native state for 16 proteins in the 18 candidates which undergo local unfolding, and consistently samples a wide range of contact ratio from native to unfolded. (Fig. S5)

As a final part of the BioEmu benchmark, we focused on the large-scale domain motions related to protein functions. LAO-binding protein (P02911) consists of two globular domains connected by three flexible peptide strands^36^. The two domains are separated by a wide cleft in the absence of a ligand, while being close to each other when binding to a ligand. The manganese-dependent inorganic pyrophosphatase (PPase, P37487) reversely adopt a closed conformation at holo state and open at apo state^37^. IDPFold2 samples a wide range of conformations that covers both states. Another case is calmodulin (P0DP23), an essential part of calcium signal transduction pathway^38^. It mainly forms an extend structure in bulk water but becomes compacted in reverse micelles^39^. See fig. S6 for all 19 cases, in which IDPFold2 predicts both reference states on 10 cases and one of the references for 8 cases, indicating the ability of IDPFold2 to identify flexible regions and the resulting motion scales.

### Fitting global and local experimental observations

While long-time MD simulations and key functional states provide references for evaluating the distribution of generated ensembles, they also introduce potential bias since they do not represent the full protein dynamics under physiological condition. Instead, experimental observations from SAXS and NMR provide experimentally derived ensemble-averaged properties and have long served as gold reference in evaluating force fields^40^. We thus used the PeptoneBench benchmark for ensemble prediction that is based on 439 SAXS profiles from SASBDB and 659 proteins with NMR chemical shifts (CS) from BMRB, advancing in both dataset scale and system diversity.

On both SAXS and CS benchmark, IDPFold2 outperformed all current deep learning approaches. To identify the disorder tendency of test proteins, PeptoneBench classifies proteins by their mean G-score ranging from 0 (fully ordered) to 1 (fully disordered). G-scores were calculated from secondary chemical shifts or predicted by ADOPT2 for those lacking experimental CS data^27^. Results indicated that IDPFold2 performed well on both benchmark dataset across order-disorder continuum. As for comparison, idpGAN and IDPFold, two previously described methods designed for IDPs specifically, showed clear advantage for IDPs compared to folded proteins on SAXS benchmark. Meanwhile, the structure prediction models AlphaFold2, ESMFold, and ESMFlow reversely performed better on folded proteins. BioEmu also maintain a balanced performance, but IDPFold2 consistently surpass these models across the whole continuum (Fig. 3A, Fig. S7). We increased the sample size of predicted ensembles and found little fluctuations in both plain and reweighted results, representing the efficiency of the independent sampling of conformations in the flow matching model. Same conclusions maintain for the CS benchmark (Fig. 3B, Fig. S8) when backmapping the generated Cα structures with cg2all^41^.

**Figure 3.**
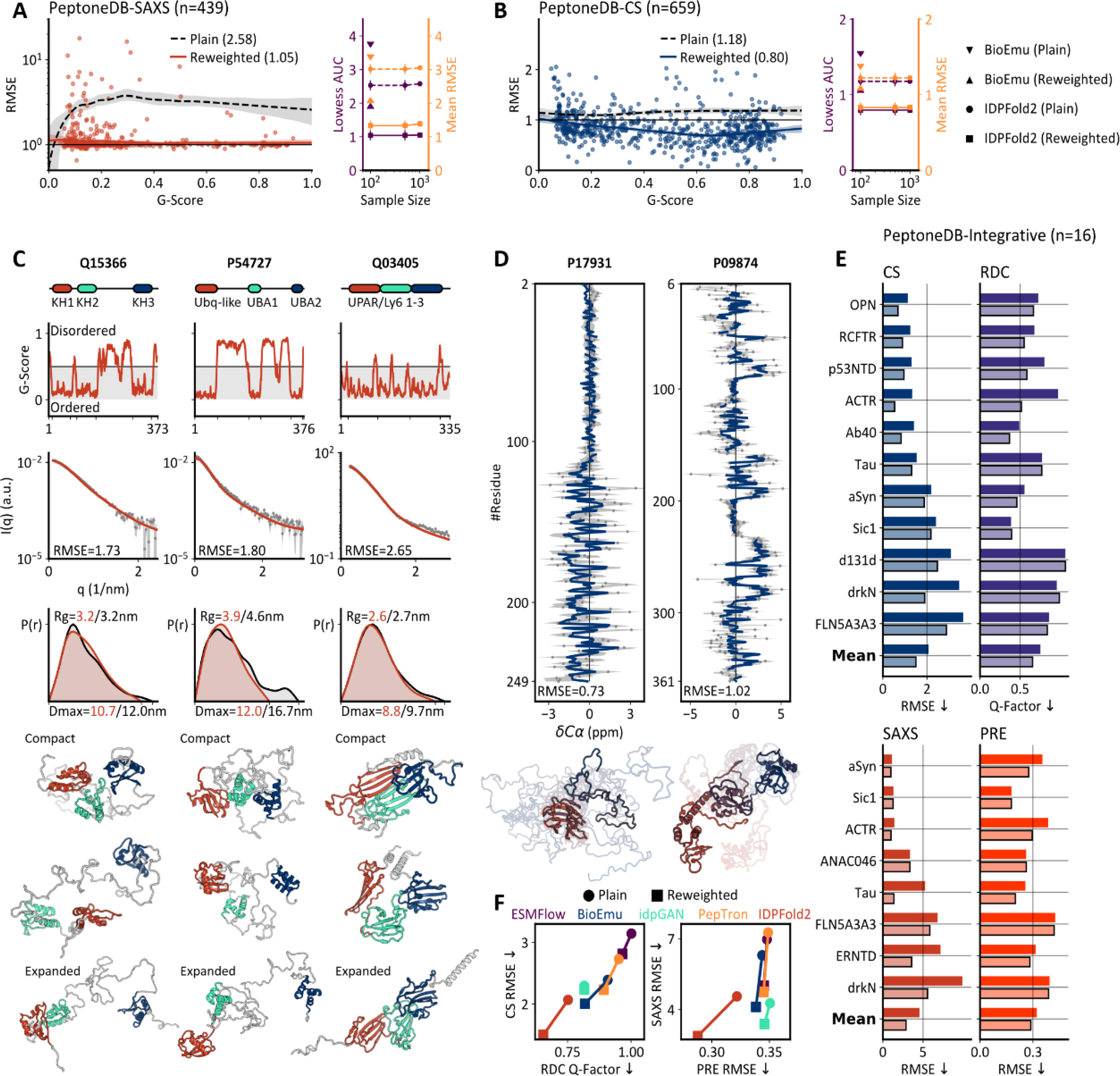
Validation against global and local experimental data. (A) SAXS benchmark. Left: Normalized RMSE vs. disorder proportion (G-score), with LOWESS regression curves. Area below LOWESS curves (AUCs) are reported. Uncertainty is estimated by bootstrapping. Right: AUC and average RMSE as a function of ensemble size. (B) CS benchmark performance (metrics same as in A). (C) Multidomain protein cases from SAXS benchmark. From top to bottom are domain assignment, SAXS profiles (intensity vs. scattering vector), pair distance distributions, and representative structures. (D) Mixed folded/disordered protein cases from CS benchmark: Top: Cα secondary chemical shifts. Bottom: generated structures. (E) Integrative benchmark. Top: Plain and reweighted RMSE on CS data, Q-factors measuring RDC agreement are reported for plain and CS-reweighted ensembles. Bottom: RMSE on SAXS and PRE data, reweighting is conducted on SAXS data. (F) Comparison of average RMSE on integrative benchmark across five generative models.

We focused specifically on the multidomain proteins in the SAXS benchmarks, whose predicted G-scores range from 0.3 to 0.5, highlighting three systems. Poly(rC)-binding protein 2 (Q15366) contains three KH domains that mediate poly(rC) binding, together contributing to negative regulation of antiviral signaling^42^. UV excision repair protein RAD23 (P54727) consists of a ubiquitin-like domain on its N-terminus acting as the tether to the proteasome, and two UBA domains that bind to cargo. It delivers ubiquitinated proteins to proteasome, thus modulating protein degradation^43^. The last case urokinase plasminogen activator receptor (uPAR, Q03405) containing three continuous repeats of UPAR/Ly6 domain. The three domains are natively arranged tightly to form a central, cone-shaped cavity, and contributes to different stages of uPA binding^44^. The three cases represent the common feature of multidomain proteins as integrated molecular bridges, where the arrangement of individual domains play an important role in their functions.

We observed close fitting of SAXS intensity curves and pair distance distributions of plain ensembles predicted by IDPFold2 (Fig. 3C). Both compact conformations and expanded ones are sampled, where the former maintained dense domain interactions while the latter arranged domains distant to each other. On RAD23 IDPFold2 sampled a low proportion of extremely expanded conformation, resulting in the underestimation of Rg. Max-entropy reweighting efficiently correlated such bias and further reduced the error as depicted in fig. 3A.

While SAXS can reflect the global compaction and domain arrangement of multidomain proteins, chemical shifts describe mainly local features like secondary structures. We here provided the secondary chemical shifts on Cα for two cases, proving that IDPFold2 had a good estimation on the secondary structures (Fig. 3D). Similar to SAXS benchmark, max-entropy effectively reduced the error between prediction and NMR observations.

Although reweighting significantly reduces the error on SAXS and CS benchmark, there’s risk that reweighting produces overfitted ensemble to a single observation. Therefore, we next used a small benchmark in which proteins were well characterized by SAXS, PRE, CS and RDC data to show that the reweighted IDPFold2 ensembles are not overfitted to certain observations. Here reweighting was conducted on either CS or SAXS data, and cross-validated using RDCs and PREs^27^. Results on both sets showed that reweighting reduced the error on CS and SAXS at the same time that error on RDC and PRE maintained or even reduced jointly (Fig. 3E). When compared to other deep-learning models we observed greater accuracy of IDPFold2 that—before reweighting—reached an error lower than some reweighted results produced by other methods (Fig. 3F, Fig. S9). In total, IDPFold2 presents significant improvements on all three benchmarks, demonstrating its effectiveness in producing accurate ensemble-averaged properties.

### Modelling multiple conformations for protein assemblies

We extended IDPFold2 to predict the ensembles of protein multimers by incorporating chain break and relative positional embeddings (Fig. 4A)^45^. Interface structures recorded in PDB were utilized for model training, informing the model about inter-chain contact modes. To validate the performance of IDPFold2 on protein multimers, we collected 88 hetero-multimers from SASBDB and 86 IDR complexes with resolved static structures in PDB.

**Figure 4.**
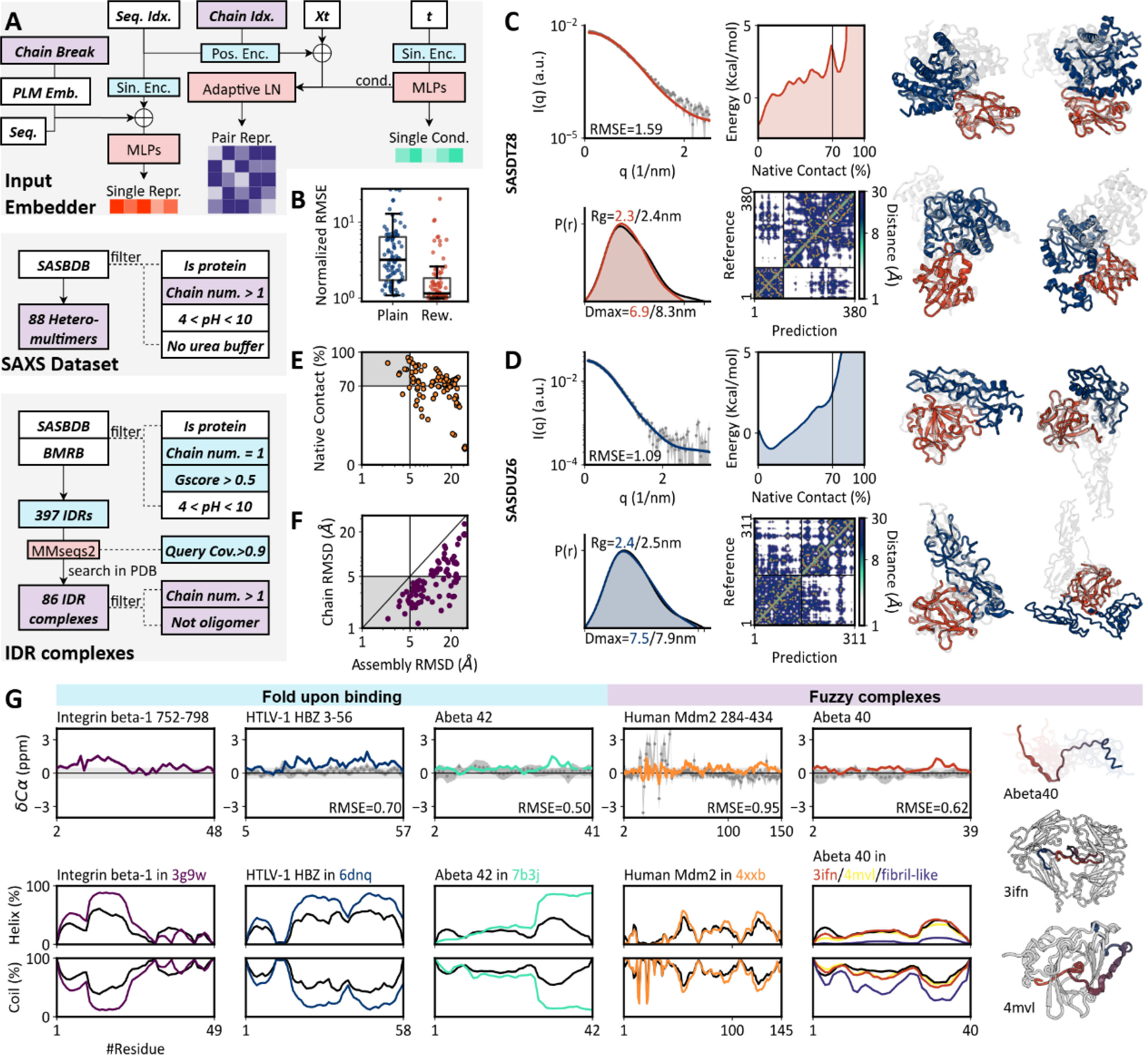
Characterizing binding modes of IDRs in assemblies. (A) Multimer modeling strategy and test dataset. (B) Normalized RMSE for SAXS profiles of hetero-multimers. (C) Analysis of SASDTZ8 assembly. Left: SAXS profile and pair distance distribution. Middle: contact fraction, contact map of reference and predicted conformation. Right: generated conformations. (D) Analysis of SASDUZ6 assembly (metrics same as in C). (E) Relationship between assembly RMSD and fraction of native contacts for IDR complexes. (F) Assembly RMSD vs. mean chain-wise RMSD. (G) IDPFold2 distinguishes IDRs that fold upon binding (e.g., Integrin beta-1) from those forming fuzzy complexes (e.g., Mdm2). Top: Cα secondary chemical shifts for monomers. Bottom: Helix and coil content in monomer (black) and multimer (colored) context.

On the SAXS dataset we calculated the normalized RMSE for intensity curves as in the PeptoneDB-SAXS monomer benchmark. IDPFold2 reached an accuracy comparable to monomer predictions after reweighting, accurately capturing the global shape of these multimers (Fig. 4B). However, SAXS data only provides little evidence on whether the predicted interfaces are correct, so we take two cases with solved structures for detailed analysis.

SASDTZ8 describes a binary multimer formed by the protein-ADP-ribose hydrolase (SpyMacro) and the glycine cleavage system H-like protein (SpyGcvH-L)^46^. Interactions mainly take place between the Zinc-dependent macrodomain in SpyMacro and the SpyGcvH-L as carrier protein, contributing to the reversal of ADP-ribosylation (Fig. 4C). IDPFold2 predicted the SAXS profile and pair distance distribution well, but only sampled a small proportion of native conformations as shown in fraction of native contacts, opposite to our observation on local-unfolding monomers. Similar results were obtained for SASDUZ6 which described the binding of domain CCP2/3 in alarmin release inhibitor and interleukin-33 (Fig. 4D)^47^. Apart from the correct interface, IDPFold2 also sampled a wide range of conformations where interfaces were partially or completely different, denoting the model still fails to estimate the exact stability of PPI.

As for the IDR multimers gathered from PDB, we also calculated the fraction of native interchain contacts preserved in generated conformations. IDPFold2 produced most native contacts for 48 in the 86 systems, among which only 7 predictions covered the exact native structure with assembly RMSD < 5Å (Fig. 4E). To check whether the high assembly RMSDs stem from interface or from the dynamics of monomers, we further calculated RMSDs for each chain. On 48 systems we got mean chain RMSD < 5Å, denoting that on this set large assembly RMSD mainly stem from biased interface. For the other 38 systems large assemble RMSD stem from either high monomer diversity or prediction error (Fig. 4F). In total, IDPFold2 was able to sample interfaces, but the scarcity of multimer data hindered the model from accurately estimating the stability of interfaces.

### Prediction conformational changes in IDR-binding

It can be difficult to study and predict the conformational ensembles of complexes between IDRs and folded domains as they often display substantial flexibility even at their binding state. While some IDRs fold into stable structures upon binding, many of the others form fuzzy complexes that retain significant structural flexibility. We evaluated IDPFold2 on six cases where the monomeric states are confirmed to be disordered by Cα chemical shifts (Fig. 4G).

IDPFold2 accurately captured transitions from disorder to ordered structures. For the Integrin beta-1 cytoplasmic tail, the model correctly predicted the transition into a stable helix upon binding the talin-2 F3 domain^48^. Similarly, for the viral protein HBZ, IDPFold2 reproduced the formation of the long helix required to wrap around the target KIX domain^49^. Crucially, the model differentiated the scale of these transitions, distinguishing the short signaling helix of Integrin from the extensive inhibitory interface of HBZ. Meanwhile, the global disorder of IDRs was preserved in fuzzy complexes. For the E3 ubiquitin ligase Mdm2 binding to ribosomal protein L11, the model maintained the requisite disordered character of the RNA-mimicking interface while correctly predicting the local formation of the essential Zinc-finger motif^50^.

IDPFold2 also resolved context-dependent behaviors in amyloid beta peptides: it predicted the helical folding of Abeta 42 when bound to the therapeutic peptide D3, versus the persistent disorder of Abeta 40 in antibody/lipocalin complexes^51–53^. For a decameric Abeta 40 assembly, IDPFold2 generated fibril-like structures as have reported in previous studies, further validating its ability to generalize across distinct IDR binding modes^54,55^.

## Discussion

We have presented IDPFold2, a generative framework that resolves a longstanding bottleneck in structural biology: the unified modeling of folded domains and IDRs. By integrating MoE architecture, the model effectively learns distinct thermodynamic signatures for different structural contexts, enabling accurate ensemble prediction for multidomain proteins and assemblies. Across benchmarks spanning static structures, MD simulations, and ensemble-averaged observables (NMR and SAXS), IDPFold2 outperforms state-of-the-art methods, offering a robust tool for exploring the dynamic proteome.

A key design choice in IDPFold2 is the use of Cα representation, which brings higher diversity in generated conformations and avoids potential chirality problems reported in previous works^45,56^. While atomistic details are indispensable for functional analysis, we demonstrate that a simple geometric backmapping via cg2all recovers local features with high fidelity, as evidenced by the accurate prediction of NMR chemical shifts. Thus, the generate-backmap protocol represents an optimal trade-off, maximizing sampling diversity and computational efficiency while hardly sacrificing local accuracy.

One limitation of IDPFold2 is the absence of kinetic pathways in generated conformational ensembles. Unlike MD simulations which derive time-dependent trajectories through bottom-up integration of energy functions, IDPFold2 generates independent equilibrium snapshots. Consequently, the ensembles lack temporal information regarding transition pathways or kinetics. There have been a series of deep-learning approaches focusing on prediction of time-dependent protein dynamics, but the limited scale and inherent uncertainty of MD simulation data have largely hindered the performance and application of these methods^57,58^.

Furthermore, while the model samples diverse conformations, the necessity for max-entropy reweighting to match experimental data suggests that the raw predicted distributions, while physically plausible, may contain residual biases. Therefore, a synergistic pipeline can be desirable: using IDPFold2 to efficiently map the conformational landscape and identify metastable states, which can then seed biased MD simulations to resolve transition barriers and kinetics^59^.

Extending generative modeling to protein assemblies reveals both the power and current limits of our approach. IDPFold2 successfully distinguishes between distinct binding modes, differentiating IDRs that undergo folding-upon-binding from those that retain fuzziness in the bound state. This distinction is critical for understanding cellular signaling, as fuzzy interactions allow for promiscuous binding and gradual affinity tuning, being distinct from the lock-and-key paradigm. We note, however, that this task remains a formidable challenge, in part perhaps because there are few methods and datasets to benchmark conformational ensembles for oligomeric assemblies. Our analysis suggests that while the model captures global assembly shapes, the scarcity of training data on multimers prevents the stabilized packing of interfaces. As observed with monomeric proteins, we anticipate that the generation of large-scale multimer simulation datasets will be required to reach higher precision for complex interfaces.

Looking ahead, the scalability of generative models remains a critical frontier. While IDPFold2 efficiently samples standard complexes, applying attention-based architectures to the megadalton-scale assemblies that govern cellular machinery is currently limited by quadratic computational costs. Overcoming this barrier will likely require the adoption of linear-complexity architectures or state-space models^60,61^. Moreover, cellular function is rarely purely proteinaceous. Expanding the chemical scope to include nucleic acids and small molecules is a necessary next step to fully capture the dynamics of regulation and metabolism, which presents a similar challenge in dataset scale and quality as in modelling protein multimers^45,62,63^.

Meanwhile, a pivotal frontier lies in decoding the regulatory layer of protein dynamics: post-translational modifications (PTMs). IDRs are evolutionary hotspots for phosphorylation, acetylation, and methylation, acting as integration hubs for cellular signaling^64,65^. Extending conditional predictions on modified residue states represents a significant step towards modeling conformational switching in response to cellular stimuli. This would provide a mechanistic bridge between genetic code and cell state, and allow researchers to simulate how specific signaling events modulate the accessibility of binding motifs or the phase behavior of condensates.

Finally, while designing IDRs and their binders holds transformative potential for synthetic biology, current methodologies predominantly focus on backbones with predefined tertiary structures. The difficulty of quantifying the dynamic properties makes IDRs challenging targets to design, with sporadic successes based on iterations of sampling methods or molecular simulations^66,67^. As a powerful structural predictor for protein ensembles, IDPFold2 possesses significant potential through two complementary avenues. First, much as the successes of protein structure prediction models in assisting designing *de novo* protein binders, IDPFold2 could provide accurate estimation of ensemble properties to computationally evaluate and filter designed IDR systems^68,69^. Secondarily, the model could serve as a plug-in design engine by inverting the generative architecture similar to the hallucination strategy, creating synthetic IDRs with precisely tuned entropic forces or specific fuzzy binding profiles^70,71^. The combined capability for validation and generative design represents an untapped engineering space that leverages the unique material properties of the dynamic proteome, paving the way for a novel computational oracle for the design of dynamic properties in proteins.

In conclusion, IDPFold2 establishes the Mixture-of-Experts framework as a versatile paradigm for modeling structural heterogeneity. By successfully integrating diverse thermodynamic regimes within a single model, it provides a flexible foundation that can be readily augmented with future algorithmic advances. This work marks a step forward in our ability to correlate structural dynamics with biological function, bringing us closer to a comprehensive, animated simulation of the living cell.

## Materials and Methods

### Dataset

The training dataset was established by integrating experimental structures and simulated conformational ensembles. The experimental component was derived from the PDB database, encompassing all entries determined by X-ray diffraction or Cryo-EM. These entries were filtered to include only structures with resolution ≤ 5.0Å. We extracted single protein chains shorter than 1,000 residues and clustered these chains at 40% sequence identity using MMSeq2, yielding 644,821 chains organized into 32,642 non-redundant clusters.

Simulated ensembles supplemented the dataset via three distinct catalogs of protein trajectories: mdCATH, IDRome-o, and AF-CALVADOS^27,72,73^. The mdCATH dataset involved 5,398 simulations of folded CATH domains, from which 100 conformations were sampled per system from each of five replicas simulated at 340K, yielding a total of 539,800 structures. IDRome-o is a set of IDR ensembles for the human proteome produced by fragment assembly using IDP-o ^27^. It contains 22,400 IDR ensembles that describe diverse conformations from fully unfold to partial fold. We collected 50 conformations for each system, totaling 1.12M structures. AF-CALVADOS features conformational ensembles of human cytosolic proteins, from which we selected 11,446 systems shorter than 1,000 residues and fetched 50 conformations per system, yielding 572,300 structures. All simulation systems were clustered at 90% sequence identity within their respective datasets.

The final training dataset was compiled by integrating the four curated sources, resulting in a total of 71,885 clusters containing approximately 2.88M structures. Structures are sampled randomly from these clusters at each training iteration. To facilitate the modeling of inter-chain interactions, binary multimers were assembled on-the-fly with a 60% probability during training. These binary combinations were pre-identified if the distance between any two Cα atoms from different chains was 10.0 Å or less. This procedure gathered 792,894 binary multimer pairs involving 559,075 chains.

As for test set, there has already been several benchmarks raised in previous studies for either folded regions or IDRs. We collected entries from the BioEmu benchmark that mainly evaluate motions of folded regions. As for IDRs, we referred to the test set provided in IDRome and CALVADOS3, which contains 73 IDRs with resolved Rg^28,74^. To provide more comprehensive evaluation, we further collected entries from BMRB and SASBDB following the protocols in Peptone-Benchmark, which contains NMR observations and SAXS profiles serving as ground truth for ensemble-averaged features.

We also collected 7 multidomain proteins with experimental Rgs from CALVADOS3 benchmark. Together with those multidomain proteins in the Peptone-Benchmark, we evaluated the performance of IDPFold2 on these proteins whose dynamics were distinct across different regions.

Finally, we established a benchmark for evaluating the predicted dynamics of protein multimers. This part mainly consisted of two compositions: the multimers resolved by SAXS and recorded in PDB. Gathering the former part followed a similar way to that used in Peptone-Benchmark, while for the latter part we searched the entries with G-score > 0.5 in Peptone-Benchmark in PDB. We finally collected 88 entries as SAXS dataset and 86 IDR complexes deposited in PDB.

Having gathered entries for test set, we removed sequences in training set of over 40% identity against test set sequences. The final training set contains 2.80M sequences belonging to 70,558 clusters, and the test set contains 1,258 monomers and 174 multimers. Details regarding the composition and construction of training and test set are provided in Table 1 and Table 2.

**Table 1.** Composition of training dataset.

| Source | Structures | Clusters |
| --- | --- | --- |
| PDB | 644,821 | 32,642 |
| PDB (multimer) | 792,894 | N/A |
| mdCATH | 539,800 | 5,387 |
| IDRome-o | 1,120,000 | 22,333 |
| AF-CALVADOS | 572,300 | 10,891 |
| Total | 2,876,921 | 71,253 |
| **Filter test set** | **2,804,053** | **70,558** |
| Total (multimers) | 729,906 | N/A |

**Table 2.** Composition of test dataset.

| Source | Systems |
| --- | --- |
| **BioEmu Benchmark** |  |
| Fast-folding proteins | 12 |
| CATH domains | 17 |
| Local unfolding | 19 |
| Domain motions | 16 |
| **Rg Benchmark** |  |
| IDRs | 73 |
| MDPs | 7 |
| **Peptone Benchmark** |  |
| SAXS | 439 |
| CS | 659 |
| Integrative | 16 |
| **Multimers** |  |
| SAXS | 88 |
| IDR complexes | 86 |
| Total | 1,258 monomers |
|  | 174 multimers |

### Flow Matching

The generation process is modeled using the flow-matching framework, a time-dependent approach that describes the transformation between a simple noise distribution and the complex data distribution of protein structures^75^. This framework defines a forward process that injects noise into the original data, and a symmetric backward process that predicts the denoised structure from Gaussian noise.

During training, the noisy structure 𝑥_𝑡_ is defined by the following formulation:

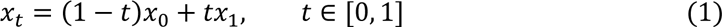

where 𝑥_0_ denotes ground truth protein structures and 𝑥_1_ is noise sampled from the standard Gaussian distribution 𝒩(0, 𝐼). The neural network parameterized by 𝜃 is trained to predict the vector field 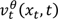 that connects 𝑥_0_ and 𝑥_1_. During inference, the conditional probability flow is modeled via an ordinary differential equation (ODE):

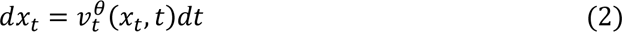

In our practice, ODE is simplified to a linear interpolation scheme for discrete-time sampling. This scheme connects each 𝑥_𝑡_ and 𝑥_𝑡−1_ at a constant interval of 0.005, which corresponds to 200 denoising steps. A single denoising step is thus given by:

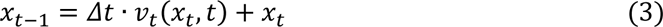

To ensure stability and facilitate model learning, all coordinate inputs and network predictions are handled at the nanometer scale.

### Model Architecture

Taking inspiration from AlphaFold3, we adopted a non-equivariant transformer architecture as the foundation of our neural network. Each transformer block operates on three primary feature sets: single representation 𝑟_𝑠_ ∈ ℝ^𝑁×𝑆^, condition representation 𝑟_𝑐_ ∈ ℝ^𝑁×𝐶^ and pair representation 𝑟_𝑝_ ∈ ℝ^𝑁×𝑁×𝑃^. Each block can be logically partitioned into a pair-biased attention module and a feed-forward network (FFN) as described by the following formulation:

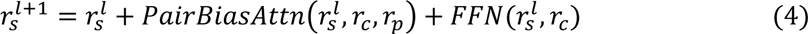

In which 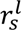 is the updated single representation at the 𝑙^𝑡ℎ^ layer. The detailed procedure of pair-biased attention and FFN are provided in Algorithm S1 and S2.

Several previous studies on protein structure generation have also employed non-equivariant transformer architectures^76–78^. However, these works primarily focused on folded proteins. Modeling the combined system of folded regions and IDRs presents a significant challenge due to the intrinsic differences in their dynamics scale and conformational behavior. We hypothesize that separately processing residues from these two distinct components, rather than forcing all residues through a single monolithic model, should yield better modeling fidelity for complex protein systems that often contain both regions.

Therefore, we introduced a Mixture-of-Experts (MoE) architecture to complement the standard forward pass of the transformer blocks, enabling the model to effectively capture the heterogeneous dynamic features within a protein. The MoE framework employs a router network that assigns tokens to different experts. This router is composed of a linear layer that takes the single representation as input and outputs softmax-weighted scores for each “expert” network. Instead of updating 𝑟_𝑠_ with a single FFN, the MoE framework updates it with a weighted-sum prediction from several FFNs (experts), which can be described by the following formulations:

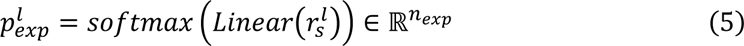

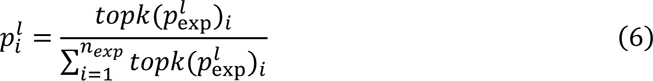

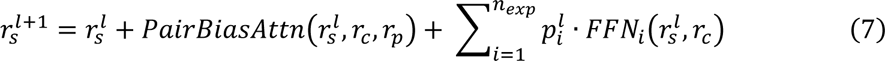

In our implementation, we utilized one shared expert and five routed experts, with two routed experts selected per token in practice. This configuration results in a total of 330M model parameters, with approximately 90M parameters activated per token.

As for model input, the initial single representation is constructed by concatenating the last-layer embedding from ESM2-650M, one-hot encoded residue types, the noisy structure coordinates, and chain-break positions. The flow-matching timestep is encoded using sine-cosine embedding and forms the condition representation. Finally, the pair representation is initialized by concatenating the pairwise distances of the noisy structure, relative positional encoding of the chain and residue indices, and the encoded timestep. Further architectural details are provided in Algorithm S3 and Algorithm S4.

### Training Strategy

We used a combination of flow-matching loss and load balancing loss within the MoE framework as the training objective, which is formally presented as:

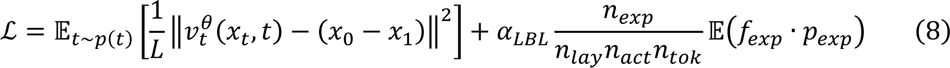

The first term represents the flow-matching loss that minimize the difference between predicted vector field 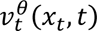 and the true velocity. 𝑝(𝑡) = 0.02𝒰(0, 1) + 0.98ℬ(1.9,1.0) denotes the timestep distribution from which t is sampled during training (𝒰 is the uniform distribution and ℬ is the Beta distribution).

The second term is the load balancing loss, which aims to ensure a balanced assignment of tokens across all experts^79^. In this term, 𝛼_𝐿𝐵𝐿_ = 0.1 denotes the weight for the load balancing loss, 𝑛_𝑒𝑥𝑝_ = 5 is the number of routed experts per layer, 𝑛_𝑙𝑎𝑦_ = 10 is the number of MoE layers, 𝑛_𝑎𝑐𝑡_ = 2 denotes activated experts per layer for each token, and 𝑛_𝑡𝑜𝑘_ is the number of tokens in one batch. 𝑓_exp_ ∈ ℝ^𝑛𝑒𝑥𝑝^ and 𝑝_𝑒𝑥𝑝_ ∈ ℝ^𝑛𝑒𝑥𝑝^ denote the number of tokens assigned to each expert and the predicted weight of each expert, respectively.

### Evaluation Metrics

To ensure rigorous validation, we compiled a comprehensive benchmark of test proteins from diverse sources, adhering to established protocols to facilitate direct comparison with prior studies.

To evaluate the fidelity of generated ensembles against Molecular Dynamics (MD) trajectories, we assessed Mean Absolute Error (MAE) and coverage of the conformational landscape projected onto the two slowest Time-lagged Independent Components (TICs). TICs were calculated on reference MD trajectories using the deeptime package, and predicted ensembles were subsequently projected onto these predefined TIC spaces. The 2D TIC space was discretized into 100*100 grids, and the free energy Δ𝐺 for each bin was derived from the occupancy probability. The metrics are defined as follows:

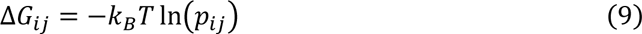

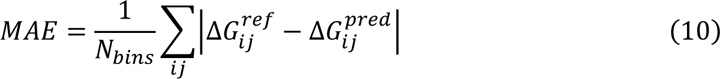

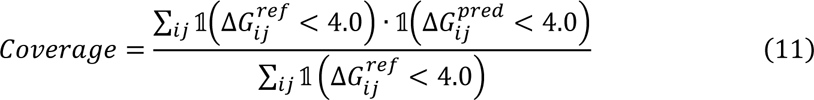

where 𝑝_𝑖𝑗_ denotes the sampled probability at the (𝑖, 𝑗)-th grid cell in the 2D TIC space, 𝑘_𝐵_ is the Boltzmann constant, and 𝑇 is the sampling temperature (set to 300 K by default). The indicator function 𝟙 equals 1 if the condition is met and 0 otherwise. Free energy values are reported in kcal/mol.

For the assessment of local unfolding and domain motions, we calculated the Root Mean Square Deviation (RMSD) against reference structures and the fraction of native contacts preserved. Native contacts were defined by a pairwise Cα distance cutoff of < 8 Å in the reference structure.

Comparison with experimental observables followed the PeptoneBench protocol. Small-angle X-ray scattering (SAXS) profiles were computed using Pepsi-SAXS, and pair distance distributions were derived using the SAXS-assistant package^80,81^. NMR chemical shifts were predicted using UCBshift^82^. Paramagnetic Relaxation Enhancement (PRE) and Residual Dipolar Couplings (RDCs) were calculated using DEERpredict and PALES, respectively^83,84^.

## Data Availability

Training and inference code is available through https://github.com/Junjie-Zhu/IDPFold2. Model checkpoints, generated ensembles and code for figures in main text are archived in https://zenodo.org/records/18239596. Supporting information in preparation.

## Acknowledgement

This work was supported by Shanghai Municipal Science and Technology Major Project, the Center for HPC at Shanghai Jiao Tong University, and the National Key Research and Development Program of China (2025YFA0921000 and 2023YFF1205102), and the National Natural Science Foundation of China (32571435). We thank Z. Zheng and Z. Fan for providing helpful discussions, and J. Yu for viewpoints on multidomain proteins.

## Notes

K.L.-L. holds stock options in, is a consultant for, and receives sponsored research from Peptone. The remaining authors declare that there is no conflict of interest.

## Reference

1. Lennicke, C. & Cochemé, H. M. Redox metabolism: ROS as specific molecular regulators of cell signaling and function. Mol. Cell 81, 3691–3707 (2021).

2. Reina-Campos, M., Scharping, N. E. & Goldrath, A. W. CD8+ T cell metabolism in infection and cancer. Nat. Rev. Immunol. 21, 718–738 (2021).

3. Choi, S. R. et al. Structural basis of microtubule-mediated signal transduction. Cell 10.1016/j.cell.2025.11.011 (2025) doi:10.1016/j.cell.2025.11.011.

4. Lai, Q. et al. Structural basis of glucose-6-phosphate transport by human SLC37A2. Nat. Struct. Mol. Biol. 1–11 (2025) doi:10.1038/s41594-025-01712-4.

5. van der Lee, R. et al. Classification of Intrinsically Disordered Regions and Proteins. Chem. Rev. 114, 6589–6631 (2014).

6. Wright, P. E. & Dyson, H. J. Intrinsically unstructured proteins: re-assessing the protein structure-function paradigm. J. Mol. Biol. 293, 321–331 (1999).

7. Holehouse, A. S. & Kragelund, B. B. The molecular basis for cellular function of intrinsically disordered protein regions. Nat. Rev. Mol. Cell Biol. 25, 187–211 (2024).

8. Wright, P. E. & Dyson, H. J. Intrinsically disordered proteins in cellular signalling and regulation. Nat. Rev. Mol. Cell Biol. 16, 18–29 (2015).

9. Beier, A. et al. Modulation of Correlated Segment Fluctuations in IDPs upon Complex Formation as an Allosteric Regulatory Mechanism. J. Mol. Biol. 430, 2439–2452 (2018).

10. Fung, H. Y. J., Birol, M. & Rhoades, E. IDPs in macromolecular complexes: the roles of multivalent interactions in diverse assemblies. Curr. Opin. Struct. Biol. 49, 36–43 (2018).

11. Sun, Q. et al. Computer-Aided Drug Discovery for Undruggable Targets. Chem. Rev. 125, 6309–6365 (2025).

12. Berman, H. M. et al. The Protein Data Bank. 10.1093/nar/28.1.235.

13. Zadorozhnyi, R., Gronenborn, A. M. & Polenova, T. Integrative approaches for characterizing protein dynamics: NMR, CryoEM, and computer simulations. Curr. Opin. Struct. Biol. 84, 102736 (2024).

14. Nettels, D. et al. Single-molecule FRET for probing nanoscale biomolecular dynamics. Nat. Rev. Phys. 6, 587–605 (2024).

15. Mugnai, M. L. et al. Sizes, conformational fluctuations, and SAXS profiles for intrinsically disordered proteins. Protein Sci. 34, e70067 (2025).

16. Su, B., Huang, K., Peng, Z., Amunts, A. & Yang, J. CryoAtom improves model building for cryo-EM. Nat. Struct. Mol. Biol. 1–11 (2025) doi:10.1038/s41594-025-01713-3.

17. Maier, J. A. et al. ff14SB: Improving the Accuracy of Protein Side Chain and Backbone Parameters from ff99SB. J. Chem. Theory Comput. 11, 3696–3713 (2015).

18. Robustelli, P., Piana, S. & Shaw, D. E. Developing a molecular dynamics force field for both folded and disordered protein states. Proc. Natl. Acad. Sci. 115, E4758–E4766 (2018).

19. Zhang, Y., Liu, H., Yang, S., Luo, R. & Chen, H.-F. Well-Balanced Force Field ff03CMAP for Folded and Disordered Proteins. J. Chem. Theory Comput. 15, 6769–6780 (2019).

20. Tesei, G. & Lindorff-Larsen, K. Improved predictions of phase behaviour of intrinsically disordered proteins by tuning the interaction range. Open Res. Eur. 2, 94 (2023).

21. Joseph, J. A. et al. Physics-driven coarse-grained model for biomolecular phase separation with near-quantitative accuracy. Nat. Comput. Sci. 1, 732–743 (2021).

22. Zheng, S. et al. Predicting equilibrium distributions for molecular systems with deep learning. Nat. Mach. Intell. 6, 558–567 (2024).

23. Volk, A. A. et al. AlphaFlow: autonomous discovery and optimization of multi-step chemistry using a self-driven fluidic lab guided by reinforcement learning. Nat. Commun. 14, 1403 (2023).

24. Lewis, S. et al. Scalable emulation of protein equilibrium ensembles with generative deep learning. Science 389, eadv9817 (2025).

25. Zhu, J. et al. Accurate Generation of Conformational Ensembles for Intrinsically Disordered Proteins with IDPFold. Adv. Sci. n/a, e11636.

26. Zhang, O., Liu, Z. H., Forman-Kay, J. D. & Head-Gordon, T. Deep Learning of Proteins with Local and Global Regions of Disorder. Preprint at 10.48550/arXiv.2502.11326 (2025).

27. Invernizzi, M. et al. Advancing Protein Ensemble Predictions Across the Order–Disorder Continuum. 2025.10.18.680935 Preprint at 10.1101/2025.10.18.680935 (2025).

28. Cao, F., von Bülow, S., Tesei, G. & Lindorff-Larsen, K. A coarse-grained model for disordered and multi-domain proteins. Protein Sci. 33, e5172 (2024).

29. Janson, G., Valdes-Garcia, G., Heo, L. & Feig, M. Direct generation of protein conformational ensembles via machine learning. Nat. Commun. 14, 774 (2023).

30. Pajkos, M., Clerc, I., Zanon, C., Bernadó, P. & Cortés, J. AFflecto: A web server to generate conformational ensembles of flexible proteins from AlphaFold models. J. Mol. Biol. 437, 169003 (2025).

31. Schnapka, V., Morozova, T. I., Sen, S. & Bonomi, M. Atomic resolution ensembles of intrinsically disordered proteins with Alphafold. 2025.06.18.660298 Preprint at 10.1101/2025.06.18.660298 (2025).

32. Lindorff-Larsen, K., Piana, S., Dror, R. O. & Shaw, D. E. How Fast-Folding Proteins Fold. Science 334, 517–520 (2011).

33. Ishikawa, Y. et al. Identification and characterization of novel components of a Ca2+/calmodulin-dependent protein kinase cascade in HeLa cells. FEBS Lett. 550, 57–63 (2003).

34. Quilliam, L. A. et al. Isolation of a NCK-associated Kinase, PRK2, an SH3-binding Protein and Potential Effector of Rho Protein Signaling *. J. Biol. Chem. 271, 28772–28776 (1996).

35. McCombe, C. L. et al. A rust-fungus Nudix hydrolase effector decaps mRNA in vitro and interferes with plant immune pathways. 10.1111/nph.18727 doi:10.1111/nph.18727.

36. Higgins, C. F. & Ames, G. F. Two periplasmic transport proteins which interact with a common membrane receptor show extensive homology: complete nucleotide sequences. Proc. Natl. Acad. Sci. 78, 6038–6042 (1981).

37. Ahn, S. et al. The “open” and “closed” structures of the type-C inorganic pyrophosphatases from *Bacillus subtilis* and *Streptococcus gordonii*1. J. Mol. Biol. 313, 797–811 (2001).

38. Tsang, W. Y. et al. CP110 Cooperates with Two Calcium-binding Proteins to Regulate Cytokinesis and Genome Stability. Mol. Biol. Cell 17, 3423–3434 (2006).

39. Xu, G., Cheng, K., Wu, Q., Liu, M. & Li, C. Confinement Alters the Structure and Function of Calmodulin. Angew. Chem. Int. Ed. 56, 530–534 (2017).

40. Cavender, C. E. et al. Structure-Based Experimental Datasets for Benchmarking Protein Simulation Force Fields [Article v0.1]. ArXiv arXiv:2303.11056v2 (2025).

41. Heo, L. & Feig, M. One bead per residue can describe all-atom protein structures. Structure 32, 97–111.e6 (2024).

42. Walter, B. L., Parsley, T. B., Ehrenfeld, E. & Semler, B. L. Distinct Poly(rC) Binding Protein KH Domain Determinants for Poliovirus Translation Initiation and Viral RNA Replication. J. Virol. 76, 12008–12022 (2002).

43. Katiyar, S., Li, G. & Lennarz, W. J. A complex between peptide:N-glycanase and two proteasome-linked proteins suggests a mechanism for the degradation of misfolded glycoproteins. Proc. Natl. Acad. Sci. 101, 13774–13779 (2004).

44. Xu, X. et al. Crystal Structure of the Urokinase Receptor in a Ligand-Free Form. J. Mol. Biol. 416, 629–641 (2012).

45. Abramson, J. et al. Accurate structure prediction of biomolecular interactions with AlphaFold 3. Nature 630, 493–500 (2024).

46. Ariza, A. et al. Evolutionary and molecular basis of ADP-ribosylation reversal by zinc-dependent macrodomains. J. Biol. Chem. 300, (2024).

47. Jamwal, A., Colomb, F., McSorley, H. J. & Higgins, M. K. Structural basis for IL-33 recognition and its antagonism by the helminth effector protein HpARI2. Nat. Commun. 15, 5226 (2024).

48. Anthis, N. J. et al. The structure of an integrin/talin complex reveals the basis of inside-out signal transduction. EMBO J. 28, 3623–3632 (2009).

49. Yang, K. et al. Structural basis for cooperative regulation of KIX-mediated transcription pathways by the HTLV-1 HBZ activation domain. Proc. Natl. Acad. Sci. 115, 10040–10045 (2018).

50. Zheng, J. et al. Structure of human MDM2 complexed with RPL11 reveals the molecular basis of p53 activation. Genes Dev. 29, 1524–1534 (2015).

51. Bocharov, E. V. et al. All-d-Enantiomeric Peptide D3 Designed for Alzheimer’s Disease Treatment Dynamically Interacts with Membrane-Bound Amyloid-β Precursors. J. Med. Chem. 64, 16464–16479 (2021).

52. Basi, G. S. et al. Structural Correlates of Antibodies Associated with Acute Reversal of Amyloid β-related Behavioral Deficits in a Mouse Model of Alzheimer Disease. J. Biol. Chem. 285, 3417–3427 (2010).

53. Rauth, S. et al. High-affinity Anticalins with aggregation-blocking activity directed against the Alzheimer β-amyloid peptide. Biochem. J. 473, 1563–1578 (2016).

54. Han, M. et al. Anle138b binds predominantly to the central cavity in lipidic Aβ₄₀ fibrils and modulates fibril formation. Nat. Commun. 16, 8850 (2025).

55. Tehrani, M. J. et al. E22G Aβ40 fibril structure and kinetics illuminate how Aβ40 rather than Aβ42 triggers familial Alzheimer’s. Nat. Commun. 15, 7045 (2024).

56. Xu, S. et al. Benchmarking all-atom biomolecular structure prediction with FoldBench. Nat. Commun. 10.1038/s41467-025-67127-3 (2025) doi:10.1038/s41467-025-67127-3.

57. Jing, B., Stärk, H., Jaakkola, T. & Berger, B. Generative Modeling of Molecular Dynamics Trajectories. Preprint at 10.48550/arXiv.2409.17808 (2024).

58. Fan, Z., Zhu, J. & Chen, H.-F. DynaFold: A Latent Diffusion Based Generative Framework for Protein Dynamic Trajectory. 2025.09.14.676071 Preprint at 10.1101/2025.09.14.676071 (2025).

59. Bhakat, S. Accelerated sampling of protein dynamics using BioEmu augmented molecular simulation. 2026.01.07.698041 Preprint at 10.64898/2026.01.07.698041 (2026).

60. Zhou, Y., et al. SeedFold: Scaling Biomolecular Structure Prediction. Preprint at 10.48550/arXiv.2512.24354 (2025).

61. Zhang, T. et al. Point Cloud Mamba: Point Cloud Learning via State Space Model. Proc. AAAI Conf. Artif. Intell. 39, 10121–10130 (2025).

62. Zhang, Y. et al. Improving diffusion-based protein backbone generation with global-geometry-aware latent encoding. Nat. Mach. Intell. 7, 1104–1118 (2025).

63. Corso, G., Stärk, H., Jing, B., Barzilay, R. & Jaakkola, T. DiffDock: Diffusion Steps, Twists, and Turns for Molecular Docking. Preprint at 10.48550/arXiv.2210.01776 (2023).

64. Bah, A. & Forman-Kay, J. D. Modulation of Intrinsically Disordered Protein Function by Post-translational Modifications. J. Biol. Chem. 291, 6696–6705 (2016).

65. Devi, B., Nag, N., Uversky, V. N. & Tripathi, T. Conditional disorder in proteins: functional transitions between order and disorder. Chem. Commun. 61, 16512–16528 (2025).

66. Pesce, F. et al. Design of intrinsically disordered protein variants with diverse structural properties. Sci. Adv. 10, eadm9926 (2024).

67. Krueger, R. K., Brenner, M. P. & Shrinivas, K. Generalized design of sequence–ensemble–function relationships for intrinsically disordered proteins. Nat. Comput. Sci. 1–12 (2025) doi:10.1038/s43588-025-00881-y.

68. Bennett, N. R. et al. Improving de novo protein binder design with deep learning. Nat. Commun. 14, 2625 (2023).

69. Pacesa, M. et al. One-shot design of functional protein binders with BindCraft. Nature 646, 483–492 (2025).

70. Frank, C. et al. Scalable protein design using optimization in a relaxed sequence space. Science 386, 439–445 (2024).

71. Zhang, B. et al. Protein language model supervised motif-scaffolding design with GPDL. Int. J. Biol. Macromol. 331, 148441 (2025).

72. Mirarchi, A., Giorgino, T. & De Fabritiis, G. mdCATH: A Large-Scale MD Dataset for Data-Driven Computational Biophysics. Sci. Data 11, 1299 (2024).

73. Bülow, S. von, Johansson, K. E. & Lindorff-Larsen, K. AF-CALVADOS: AlphaFold-guided simulations of multi-domain proteins at the proteome level. 2025.10.19.683306 Preprint at 10.1101/2025.10.19.683306 (2025).

74. Tesei, G. et al. Conformational ensembles of the human intrinsically disordered proteome. Nature 626, 897–904 (2024).

75. Song, Y., Durkan, C., Murray, I. & Ermon, S. Maximum Likelihood Training of Score-Based Diffusion Models. Preprint at 10.48550/arXiv.2101.09258 (2021).

76. Qu, W. et al. P(all-atom) Is Unlocking New Path For Protein Design. 2024.08.16.608235 Preprint at 10.1101/2024.08.16.608235 (2025).

77. Geffner, T., et al. Proteina: Scaling Flow-based Protein Structure Generative Models. Preprint at 10.48550/arXiv.2503.00710 (2025).

78. Geffner, T., et al. La-Proteina: Atomistic Protein Generation via Partially Latent Flow Matching. Preprint at 10.48550/arXiv.2507.09466 (2025).

79. Dai, D., et al. DeepSeekMoE: Towards Ultimate Expert Specialization in Mixture-of-Experts Language Models. Preprint at 10.48550/arXiv.2401.06066 (2024).

80. Grudinin, S., Garkavenko, M. & Kazennov, A. Pepsi-SAXS: an adaptive method for rapid and accurate computation of small-angle X-ray scattering profiles. Acta Crystallogr. Sect. Struct. Biol. 73, 449–464 (2017).

81. Ramirez, C. et al. SAXS Assistant: Automated SAXS analysis for structural discovery in biologics and polymeric nanoparticles. Biophys. J. 124, 3772–3786 (2025).

82. Li, J., C. Bennett, K., Liu, Y., V. Martin, M. & Head-Gordon, T. Accurate prediction of chemical shifts for aqueous protein structure on “Real World” data. 10.1039/C9SC06561J (2020) doi:10.1039/C9SC06561J.

83. Tesei, G. et al. DEER-PREdict: Software for efficient calculation of spin-labeling EPR and NMR data from conformational ensembles. PLOS Comput. Biol. 17, e1008551 (2021).

84. Zweckstetter, M. NMR: prediction of molecular alignment from structure using the PALES software. Nat. Protoc. 3, 679–690 (2008).

